# Cocaine memory reactivation induces functional adaptations of fast-spiking interneurons in the rat medial prefrontal cortex

**DOI:** 10.1101/868299

**Authors:** Emily T. Jorgensen, Angela E. Gonzalez, John H. Harkness, Deborah M. Hegarty, Delta J. Burchi, Jake A. Aadland, Sue A. Aicher, Barbara A. Sorg, Travis E. Brown

**Author notes:** To whom correspondence should be addressed, Travis E. Brown, Ph.D., University of Wyoming, 1000 E University Ave., Dept 3375, Laramie, WY 82071.

## Abstract

Perineuronal nets (PNNs) are specialized extracellular matrix structures that ensheathe parvalbumin-containing fast-spiking interneurons (PV FSIs) and play a key role in neuroplasticity. We previously showed that PNNs within the prelimbic prefrontal cortex (PL PFC) are required for the maintenance of cocaine-associated memories following cocaine memory reactivation. However, how cocaine memory reactivation affects PNNs, PV, and corresponding changes in PV FSI function are unknown. In this study, we characterized the electrophysiological properties of PV FSIs and corresponding changes in PNN and PV intensity within the PL PFC prior to and after cocaine memory reactivation. Adult male Sprague-Dawley rats were trained to acquire cocaine-conditioned place preference (CPP) and, following cocaine-CPP memory reactivation (30 m, 2 h, and 24 h post-reactivation), we measured PNN intensity (determined by *Wisteria floribunda* agglutinin [WFA] staining) as well as PV intensity using immunohistochemistry. The intensity of PV staining was reduced at all time points following memory reactivation with no changes in WFA intensity. Using whole-cell electrophysiology we found a reduction in the number of action potentials at 30 m and 2 h that returned to control levels by 24 h. The attenuation in firing was accompanied by a presumed compensatory increase in excitatory synaptic transmission, which was corroborated by an increase in VGluT1 puncta apposing PV/PNN neurons. Collectively, our results indicate that cocaine memory reactivation decreases PV intensity, which may play a role in decreasing excitation of PV FSIs. Thus, the inhibitory tone onto pyramidal neurons may be decreased following memory reactivation, resulting in an increase in PFC output to promote cocaine-seeking behaviors.

## Introduction

Repeated drug exposure causes persistent drug-associated memories to form, which can contribute to relapse following abstinence [1–3]. Drug-seeking behaviors can be diminished when the drug-associated memory is weakened or disrupted during the “reconsolidation window”, a period of up to 6 h after memory reactivation [4, 5], when the memory is labile and susceptible to modification [6–9]. Notably, a memory retrieval-extinction procedure has been shown to reduce long-term craving in humans for up to 6 months, which suggests that modifying drug-associated memories via a reconsolidation process is an effective strategy to diminish drug relapse [8, 10]. However, relatively few studies have examined the neuronal changes associated with memory reactivation. In this study, we characterized the electrophysiological properties of parvalbumin (PV)-containing fast-spiking interneurons (FSIs) and corresponding changes in perineuronal net (PNN) and PV intensity within the prelimbic prefrontal cortex (PL PFC) prior to and after cocaine memory reactivation.

PV is a high-affinity calcium buffer and is a reliable identifier of GABAergic FSIs within the cortex [11–13]. PV-containing FSIs synapse directly onto layer V pyramidal neurons within the medial PFC and heavily influence excitatory output [14]. Deficits in PV cells have been implicated in multiple disorders of the nervous system, including schizophrenia and autism [15–18]. In addition, PV cells play an important role in behavioral and molecular aspects of learning and memory [19, 20]. However, to our knowledge, nothing is known about the role of PV FSIs in the development and maintenance of drug-associated memories following memory reactivation. In the PFC, PV FSIs are routinely associated with perineuronal nets (PNNs), which we have shown to mediate cocaine-associated memories after memory reactivation [21].

PNNs are specialized extracellular matrix structures composed of chondroitin sulfate proteoglycans, hyaluronin, and other proteins that appear during critical periods of development [22]. Within the PFC, PNNs primarily surround PV FSIs and broadly contribute to neuronal stability [23]. In addition, but not limited to, PNNs protect neurons from oxidative stress [24–26], impact cognitive performance and memory [27, 28], contribute to the establishment and maintenance of fear memories [29–31], and are susceptible to modification following exposure to various substances, including high-fat diets [32, 33] and cocaine [21, 34]. Recent work by us and others has shown that degradation of PNNs attenuates cocaine-conditioned place preference (cocaine-CPP) and cocaine self-administration [21, 35–37]. However, the relationship between changes in PV, PNNs, and the associated adaptations in neuronal excitability and synaptic transmission of PV FSIs within the PFC following cocaine memory reactivation is unknown.

Our previous study showed that degradation of PNNs with chondroitinase-ABC (Ch-ABC) in the PL PFC attenuated cocaine-CPP and increased the firing of pyramidal neurons within the PL PFC [21]. In addition, we observed a decrease in the frequency of miniature inhibitory events onto pyramidal neurons after PNN degradation. We hypothesized that the increase in pyramidal firing was the result of a decrease in inhibitory tone onto pyramidal neurons within the PL PFC, which would be consistent with a decreased firing rate of FSIs that has been observed within the medial nucleus of the trapezoid body after PNN degradation [38].

In the present study, we used whole-cell electrophysiology to measure intrinsic excitable properties and changes in synaptic transmission onto FSIs surrounded by PNNs, as measured by neurons that were positive for *Wisteria floribunda* agglutinin (a marker for PNNs, [WFA]) after cocaine memory reactivation. We used immunohistochemistry to determine whether cocaine memory reactivation influenced the intensity of PV and/or WFA staining. We also used immunocytochemistry and confocal microscopy to quantify GABAergic and glutamatergic puncta onto WFA^+^ PV FSIs. We found that cocaine memory reactivation transiently reduced the firing rate of WFA^+^ PV FSIs. The reduction in firing was accompanied by an increase in pre- and postsynaptic excitatory synaptic transmission and an increase in VGluT1 puncta around PV/WFA cells. PV intensity was reduced at all time points tested (from 30 min up to 24 h post-reactivation), while WFA intensity was not altered. Our results highlight that cocaine memory reactivation in the absence of cocaine significantly alters the synaptic and electrical properties of WFA^+^ FSIs and their associated proteins, which modulate pyramidal output and influences drug-seeking behavior.

## Materials and Methods

### Animals

All procedures were performed in accordance with the National Institutes of Health’s *Guidelines for the Care and Use of Laboratory Animals* and with approval from the Institutional Animal Care and Use Committee at the University of Wyoming and Washington State University. A total of 69 male Sprague Dawley rats were used for these experiments. All rats were bred in house with *ad libitum* access to food and water and weighed ~300–400 g during the experiment. Rats were housed in a temperature (25°C) and humidity-controlled room maintained on a 12 h light/dark cycle, with lights on at 0700. All efforts were made to minimize the number of animals used in the experiments and to reduce the amount of pain and suffering.

### Cocaine-Conditioned Place Preference

All CPP experiments were conducted during the light phase of the daily light/dark cycle. The CPP apparatuses consist of three Plexiglas compartments, including two primary outer chambers (28 × 21 × 21 cm), one with black walls and metal rod floors and the other with white walls and wire mesh floors. The center chamber (12 × 21 × 21 cm) has gray walls with solid gray Plexiglas flooring (Med Associates, Fairfax, VT). Side preference was automatically recorded with infrared photocell beams within the apparatus. Manual guillotine doors separated each chamber, confining the rat to one side of the CPP apparatus when necessary.

Rats received two days of 15 min exposure to all three chambers: the first day served as a habituation day and the second day served as a test for initial preference of sides of the apparatus. Time spent within each compartment was recorded. The cocaine-paired chamber was chosen by counterbalancing the preferred and non-preferred sides in addition to the black and white chambers. Saline and cocaine (gift from National Institute of Drug Abuse) were administered on alternate days for six training days. Each rat received three saline (1 mL/kg, intraperitoneal [i.p.]) and three cocaine (15 mg/kg, i.p.) pairings. Following cocaine or saline injections, each rat was confined to the assigned chamber for 25 min [21]. Following cocaine-CPP training, rats were given a memory reactivation session in the drug-free state 24 h after their last training injection by placing them in the central chamber with access to all chambers for 15 min. Rats spending more time on the cocaine-paired chamber compared to the initial preference day were considered to have attained a place preference. Animals were euthanized at select times after the memory reactivation session (30 min, 2 h, and 24 h). To ensure we measured an effect of cocaine memory reactivation and not previous cocaine training, some rats were not given a memory reactivation session but were removed directly from their home cage 24 h following cocaine-CPP training (t = 0 min).

### Whole-cell patch clamp electrophysiology

For whole-cell patch clamp, tissue was prepared as previously described [21, 34]. Following perfusion, rats were decapitated and coronal slices (300 μm) containing the PL PFC (3.2-3.7 mm from bregma, [39]) were in ice-cold recovery solution using a vibratome (Leica VT1200S). The composition of recovery solution was (in mM): 93 NMDG, 2.5 KCl, 1.2 NaH_2_PO_4_, 30 NaHCO_3_, 20 HEPES, 25 glucose, 4 sodium ascorbate, 2 thiourea, 3 sodium pyruvate, 10 MgSO_4_(H_2_O)_7_, 0.5 CaCl_2_(H_2_O_2_), and HCl added until pH was 7.3-7.4 with an osmolarity of 300-310 mOsm. Prior to recording, slices were incubated for 1 h in room temperature holding solution. Holding solution composition was (in mM): 92 NaCl, 2.5 KCl, 1.2 NaH_2_PO_4_, 30 NaHCO_3_, 20 HEPES, 25 glucose, 4 sodium ascorbate, 2 thiourea, 3 sodium pyruvate, 2 MgSO_4_(H_2_O)_7_, 2 CaCl_2_(H_2_O_2_), and 2 M NaOH added until pH reached 7.3-7.4 and osmolarity was 300-310 mOsm. Immediately before recording, slices were incubated in room-temperature holding solution containing WFA (1 μg/mL) for 5 min to stain for PNNs. The recording chamber was perfused constantly at 31.0 °C at a rate of 4-7 mL/min of aCSF. Composition of aCSF was (in mM): 119 NaCl, 2.5 KCl, 1 NaH_2_PO_4_, 26 NaHCO_3_, 11 dextrose, 1.3 MgSO_4_(H_2_O)_7_, and 2.5 CaCl_2_(H_2_O)_2_. CellSens software (Olympus) was used to identify WFA^+^ (fluorescing) neurons in layers IV and V of the PL mPFC, and later patched (Figure 1D for representation). Patching pipettes were pulled from borosilicate capillary tubing (Sutter Instruments, CA USA) and the electrode resistance was typically 4-7 mOhms. Pipettes were filled with K-Gluconate intracellular solution for intrinsic experiments. K-Gluconate composition was (in mM): 120 K-Gluconate, 6 KCl, 10 HEPES, 4 ATP-Mg, 0.3 GTP-Na, 0.1 EGTA, and KOH was added to bring pH to ~7.2. Cells were current-clamped at −70mV and 10 current steps were injected starting at −100pA and ending at 800pA. The duration of each recording was no longer than 5 min per cell with each sweep lasting 100 ms. Elicited action potentials were recorded, counted, and analyzed using pClamp10.3. (Clampfit, Axon Instruments, Sunnyvale, CA).

To record miniature inhibitory postsynaptic currents (mIPSCs) the aCSF bath contained: 6,7-dinitroquinoxaline-, 2,3-dione (DNQX; 10μM), strychnine (1μM), and tetrodotoxin (1μM) to block AMPA, glycine receptors, and sodium channels, respectively. Pipettes were filled with CsCl intracellular solution. CsCl solution composition was (in mM): 117 CsCl, 2.8 NaCl, 5 MgCl_2_, 20 HEPES, 2 Mg^2+^ATP, 0.3 Na^2+^GTP, 0.6 EGTA, and sucrose to bring osmolarity to 275-280 mOsm and pH to ~7.25. To record miniature excitatory postsynaptic currents (mEPSCs) the aCSF contained picrotoxin (100μM) and tetrodotoxin (1 μM) to block GABA receptors and sodium channels, respectively. Patch pipettes were filled with KCl intracellular solution. KCl solution composition was (in mM): 125 KCL, 2.8 NaCl, 2 MgCl_2_, 2 ATP-Na^+^, 0.3 GTP-Li^+^, 0.6 EGTA, and 10 HEPES. Cells were voltage-clamped at −70mV and input resistance and series resistance were monitored throughout experiments. Miniature recordings were 3 minutes long with 1 minute per sweep. Only one sweep per cell was used for data analysis. Criteria were met if the cell’s access resistance was below 30 MΩ. mIPSCs and mEPSCs were amplified and recorded using pClamp10.3. Mini Analysis Program Demo (Synaptosoft Inc, GA USA) was used to measure miniature amplitudes and frequencies.

**Figure 1.**
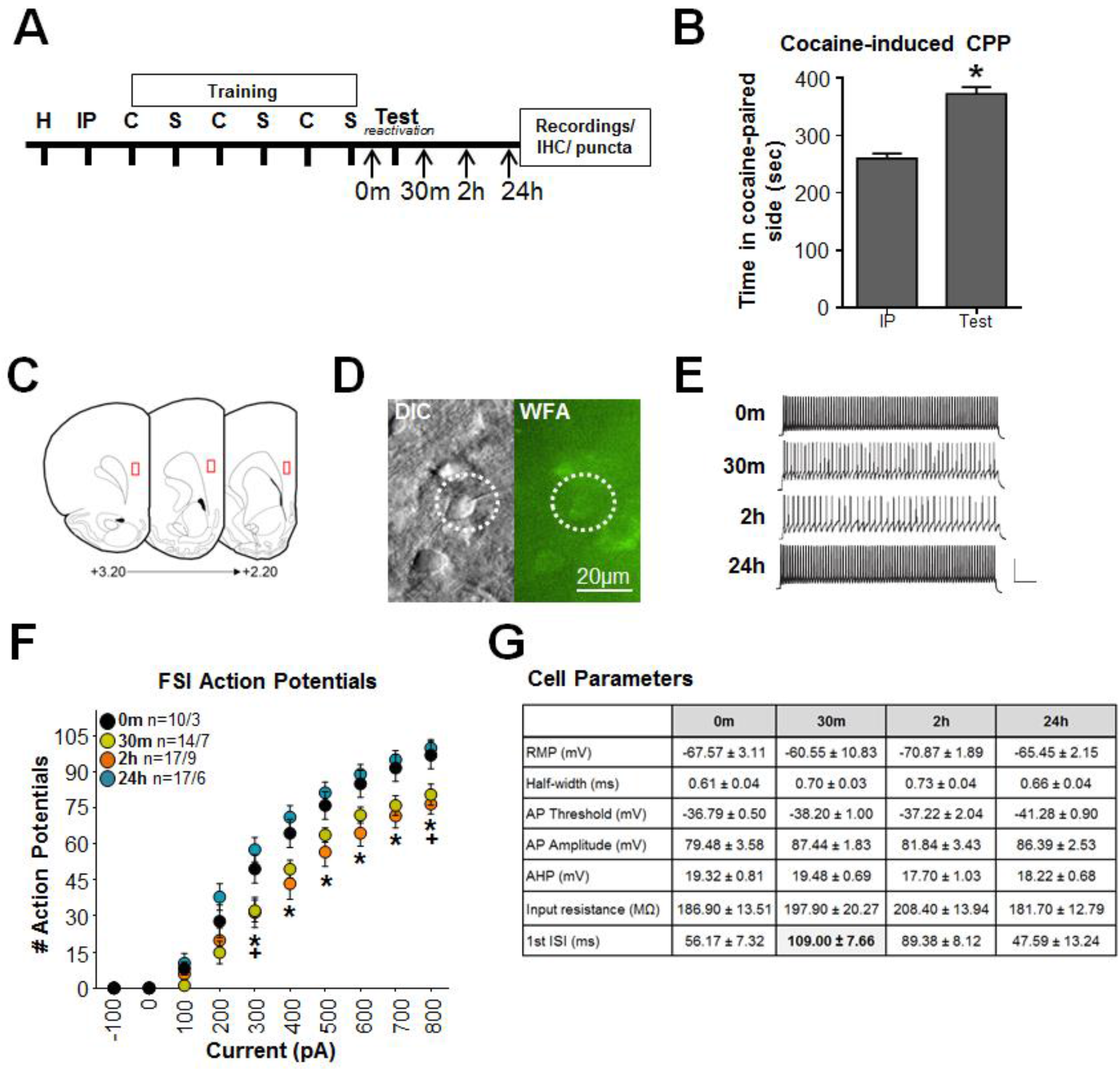
Cocaine-CPP memory reactivation decreases the firing rate of PNN surrounded FSIs within the PL mPFC. **(A)** Experimental timeline for cocaine treated animals. H=15 min habituation for CPP apparatus; IP=15 min initial preference for cocaine-paired chamber prior to training; C=25 min 15 mg/kg, intraperitoneal (i.p.) cocaine-pairing; S=25 min 1 mL/kg i.p. saline-pairing; T=test for cocaine-paired chamber after cocaine training procedure in which no cocaine was administered; arrows indicate when rats were euthanized for analysis; **(B)** Time spent in cocaine-paired chamber on initial preference and test day for CPP. Rats conditioned with cocaine (*n =* 66) show place preference; **(C)** Schematic of coronal brain sections through PFC where recordings were performed, red boxes indicates recording regions within layers IV and V of the mPFC; **(D)** DIC and WFA^+^ images of a single cell recorded in PFC; **(E)** Representative traces of action potentials evoked by 500pA current injection for cells at different times after reactivation test. Scale bar represents 50ms, 50mV; **(F)** Average numbers of action potentials across range of current injections for different groups post reactivation, as well as control groups. N-sizes reported as cell #/animal #: (t = 0 (10/3); t = 30m (14/7); t = 2h (17/9); t = 24h (17/6)). * indicate significant reduction in APs at the 30 m and 2 h time points when compared to 0 m p<0.01; **(G)** Intrinsic properties of patched neurons for different groups. 1^st^ ISI increased at 30m when compared to 0 m (p<0.05).

### Immunohistochemistry and Imaging

#### WFA and PV labeling and imaging

PV and WFA were stained as previously described [34, 40, 41]. All images (1.194 pixels/μm) were compiled into summed images using ImageJ macro plug-in Pipsqueak™ (https://labs.wsu.edu/sorg/research-resources/), scaled, and converted into 8-bit, grayscale, tiff files. Pipsqueak™ was run using the “double-label analysis” function in “semi-automatic mode” to select ROIs to identify individual PV+ cells and PNNs, which were then verified by a trained experimenter who was blinded to the experimental conditions. The plug-in compiles this analysis to identify single [41] and double-labeled neurons [40]; (https://ai.RewireNeuro.com).

### Puncta labeling and imaging

Immunohistochemical methods were like those previously described [42, 43]. The primary antibody cocktail consisted of goat anti-glutamic acid decarboxylase 65/67 (GAD 65/67, Santa Cruz Biotechnology, Cat# sc-7513, RRID: AB_2107745, 1:100), rabbit anti-parvalbumin (Novus Biologicals, Cat# NB120-11427, RRID:AB_791498, 1:1000), and guinea pig anti-vesicular glutamate transporter 1 (VGluT1; EMD Millipore, Cat# AB5905, RRID: AB_2301751, 1:5000). After 40 h primary antibody incubation, tissue sections were rinsed in TS and then incubated with a cocktail of fluorescently labeled secondary antibodies for 2 h, light-protected, at room temperature. The secondary antibody cocktail consisted of Alexa Fluor 488 donkey anti-goat (ThermoFisher Scientific, Cat# A11055, RRID: AB_2534102, 1:800), Alexa Fluor 546 donkey anti-rabbit (ThermoFisher Scientific, Cat# A10040, RRID: AB_2534016, 1:800;); and Alexa Fluor 647 donkey anti-guinea pig (Jackson ImmunoResearch Laboratories, Cat# 706-605-148, RRID: AB_2340476, 1:800). Tissue sections were rinsed again in TS and then incubated in biotinylated WFA (Vector Laboratories, Cat# B-1355, RRID: AB_2336874, 1:50) for 2 h at room temperature. Following TS rinses, tissue sections were incubated for 3 h at room temperature in Alexa Fluor 405-conjugated streptavidin (ThermoFisher Scientific, Cat# S32351, 6.25 μg/ml). Finally, tissue sections were rinsed in TS followed by PB before being mounted with 0.05 M PB onto gelatin-coated slides to dry. Slides were coverslipped with Prolong Gold Antifade Mountant (ThermoFisher Scientific) and light-protected until imaging. Anatomical landmarks were used to determine representative caudal and rostral sections of prelimbic cortex that were within +3.5 to +4.2 mm from Bregma [39]. Two high magnification images were taken at each level of the prelimbic cortex (2 images/level × 2 levels/animal = 4 images/animal). Images were captured on a Zeiss LSM 780 confocal microscope with a 63 × 1.4 NA Plan-Apochromat objective (Carl Zeiss MicroImaging1024 pixel resolution). Optical sectioning produced Z-stacks bounded by the extent of fluorescent immunolabeling throughout the thickness of each section. Using Zen software (Carl Zeiss, RRID SCR_013672), PV neurons in each confocal stack were identified and assessed for the presence of a nucleus and whether the entire neuron was within the boundaries of the field of view; only these PV neurons were included in the analysis. The optical slice through the nucleus at which the ellipsoidal minor axis length of each PV neuron reached its maximum was determined. This length was recorded as the cell diameter for each PV neuron. A Z-stack of that optical slice plus one optical slice above and one below was created resulting in a 1.15 μm Z-stack through the middle of each PV neuron; these subset Z-stacks were used for puncta apposition analysis.

For puncta analysis, the presence of WFA labeling in close proximity to the PV neuron surface was assessed for each PV neuron. A PV neuron was considered to have a PNN if there was any WFA labeling around any part of the PV neuron surface as seen by the observer. GAD65/67 and VGluT1-labeled puncta were then assessed separately using the *Spots* segmentation tool as previously described [34]. Image analysis of GABAergic and glutamatergic appositions onto PV-labeled neurons was performed using Imaris 8.0 software (BitPlane USA, Concord, MA, RRID: SCR_007370) on an offline workstation in the Advanced Light Microscopy Core at Oregon Health & Science University by a blinded observer [42, 43].

### Analysis

All statistical tests were conducted using Prism 6 (GraphPad Software). For behavioral experiments, we compared the initial preference and post-training test using a paired t-test. For intrinsic electrophysiological analysis, data were analyzed with a two-way RM ANOVA followed by a Dunnett’s multiple comparison test to compare significance from non-reactivation (t = 0 min). For intrinsic properties, data were analyzed with a one-way ANOVA followed by a Tukey’s multiple comparison test to compare significance from t=0. For miniature electrophysiological analysis, data were analyzed with a Kolmogorov-Smirnov test when assessing cumulative probability distributions and a one-way ANOVA followed by a Tukey’s multiple comparison test when assessing average miniature data. For WFA and PV intensity, control group mean cell intensities were used to calculate normalized intensities for each label, respectively. Distributions of normalized intensities were then compared between experimental groups, within stain type, using a Kruskal Wallis test to assess changes in the distribution of intensities among groups. Comparison among treatment groups for the number of PV- and WFA – labeled cells was done using a one-way ANOVA. For puncta analysis, data were analyzed using a Kruskal-Wallis test followed by a Dunn’s multiple comparison test. All results are summarized as mean ± standard error of the mean (SEM). Significance was considered at p < 0.05.

## Results

### Cocaine memory reactivation alters excitability of FSIs

Figure 1A shows the experimental timeline. There was a significant increase in place preference for the cocaine-paired chamber during the drug-free CPP test (Figure 1B, Test/memory reactivation session) compared to the initial preference (IP) day in all rats (IP = 252 ± 9 seconds, Test = 373 ± 11 seconds; t(57) = 8.88, p<0.01). There were no significant differences within each experimental group (rats used for electrophysiology, immunohistochemistry, and puncta analysis, data not shown), hence, behavioral data were pooled.

Memory reactivation attenuated the number of current-induced action potentials in WFA^+^ FSIs compared to the non-reactivated rats, this attenuation was present at both 30 min and 2 h following memory reactivation and returned to control levels by 24 h (Figure 1F; reactivation time points: F_(3,54)_= 6.69, p<0.01; current step: F_(9,486)_= 568.6, p<0.01; interaction: F_(27,486)_= 3.98, p<0.01). Within the intrinsic properties, the first ISI was increased 30 min following memory reactivation compared to t=0 (Figure 1G; F_(3,54)_= 7.98, p<0.01).

### Cocaine memory reactivation alters synaptic transmission onto FSIs

Memory reactivation increased average mEPSC frequencies 30 min, 2 h, and 24 h after memory reactivation compared to non-reactivated controls (Figure 2A; 0 min = 0.84Hz, 30 min = 4.06Hz, 2 h = 2.67, and 24 h = 3.38; F_(3,77)_= 10.04, p<0.01). The average miniature data is representative of collapsed events for each cell to get a sense of individual cell variability. We also provide cumulative data of all the miniature events; this is more sensitive and representative of all miniature events. Hence, we can detect significance in the cumulative plots that are trends when the data were collapsed to the average data. Additionally, we found differences in the cumulative distributions of both mEPSC inter-event intervals and amplitudes. Compared to t = 0, the inter-event intervals of all other time-points shifted leftward, indicating that the inter-event interval was smaller (Figure 2B), which indicates an increased frequency. There was also a potentiation of the mEPSC amplitude 30 min after memory reactivation compared to non-reactivated controls (Figure 2C; 0 min = 13.24pA, 30 min = 19.21pA, 2 h = 15.99pA, and 24 h = 14.66pA; F_(3,77)_= 10.97, p<0.01). In addition, there was a rightward shift in the cumulative distribution of individual amplitudes (Figure 2D), indicating a potentiation in amplitudes at all time points when compared to t = 0 (p<0.01). There were no significant differences in average mIPSC frequency (Figure 2E) or amplitude (Figure 2G). However, we did observe significant shifts in the cumulative distributions of the mIPSC inter-event intervals (Figure 2F) and individual amplitudes (Figure 2H);209 inter-event intervals shifted right from t = 0, where amplitudes shifted left from t = 0 (p<0.01), indicative of a decrease in inhibitory synaptic transmission.

**Figure 2:**
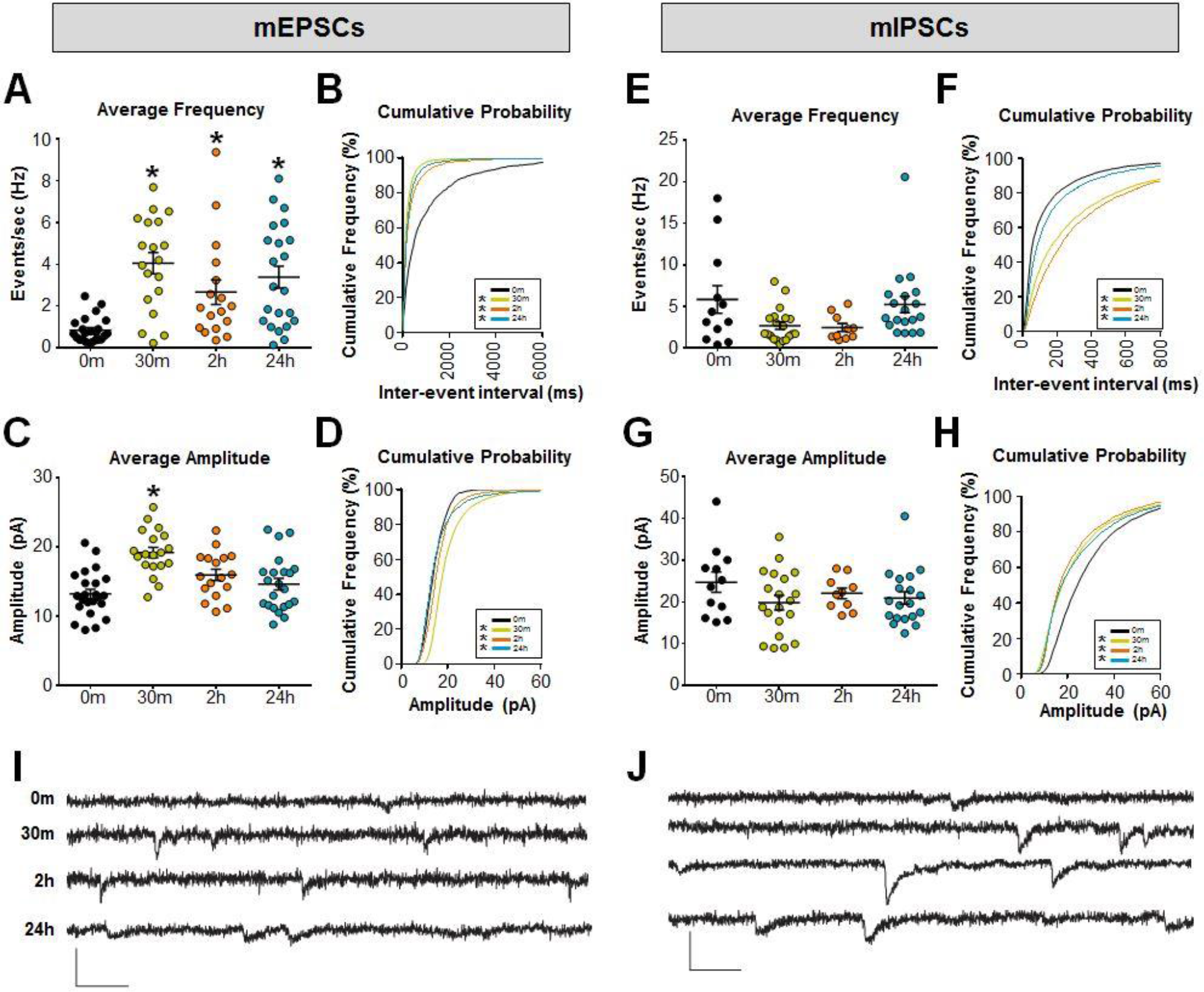
Cocaine-CPP memory reactivation increases mEPSC amplitude and frequency of PNN surrounded FS IN within the PL PFC. (mEPSCs) Cocaine-CPP memory reactivation significantly increased glutamatergic synaptic transmission. **(A)** Average frequency (Hz) of mEPSC events/neuron. N-sizes reported as cell #/animal #: (t = 0 (23/4); t = 30m (19/3); t = 2 h (17/3); t = 24 h (22/3)); * p < 0.05, compared with t=0 time point. **(B)** Cumulative probability of inter-event intervals for all events; *p<0.001, all time points relative to t=0. **(C)** Average amplitudes (pA) of mEPSC events/neuron; * p < 0.05, compared with t=0 time point; and **(D)** cumulative probability of amplitudes for all events (left); *p<0.001, t=0 compared to all time points. **(mIPSCs)** Cocaine-memory reactivation significantly altered cumulative distributions of inhibitory synaptic transmission. **(E)** Average frequency of mIPSCs events/neuron. N-sizes: (t=0 (12/3); t= 30 m (20/3); t = 2 h (10/3); t = 24 h (19/3)); **(F)** Cumulative probability of inter-event intervals for all events (right); *p<0.001, t=0 compared to all time points. **(G)** Average amplitudes of mIPSCs events/neuron and; **(H)** cumulative probability of amplitudes for all events (right); *p<0.001, all time points compared to t=0. **(I)** Representative traces from mEPSCs; **(J)** Representative traces from mIPSCs. Scale bar represents 50pA, 500μs.

### Cocaine-conditioned place preference alters intensity of PNNs and PV of FSIs

Brains were examined for PV and WFA staining intensity at the times indicated in Figure 1A. Figure 3B shows that the intensity of PV in WFA-surrounded cells was decreased at all time points (30 min, 2 h, and 24 h) compared with non-reactivated controls (p<0.0001 for all groups). There was no significant difference in the intensity of WFA around PV cells 30 m, 2 h, or 24 h after reactivation (Figure 3C). There was also no difference in the number of PV/WFA double-labeled neurons across time points (data not shown).

**Figure 3.**
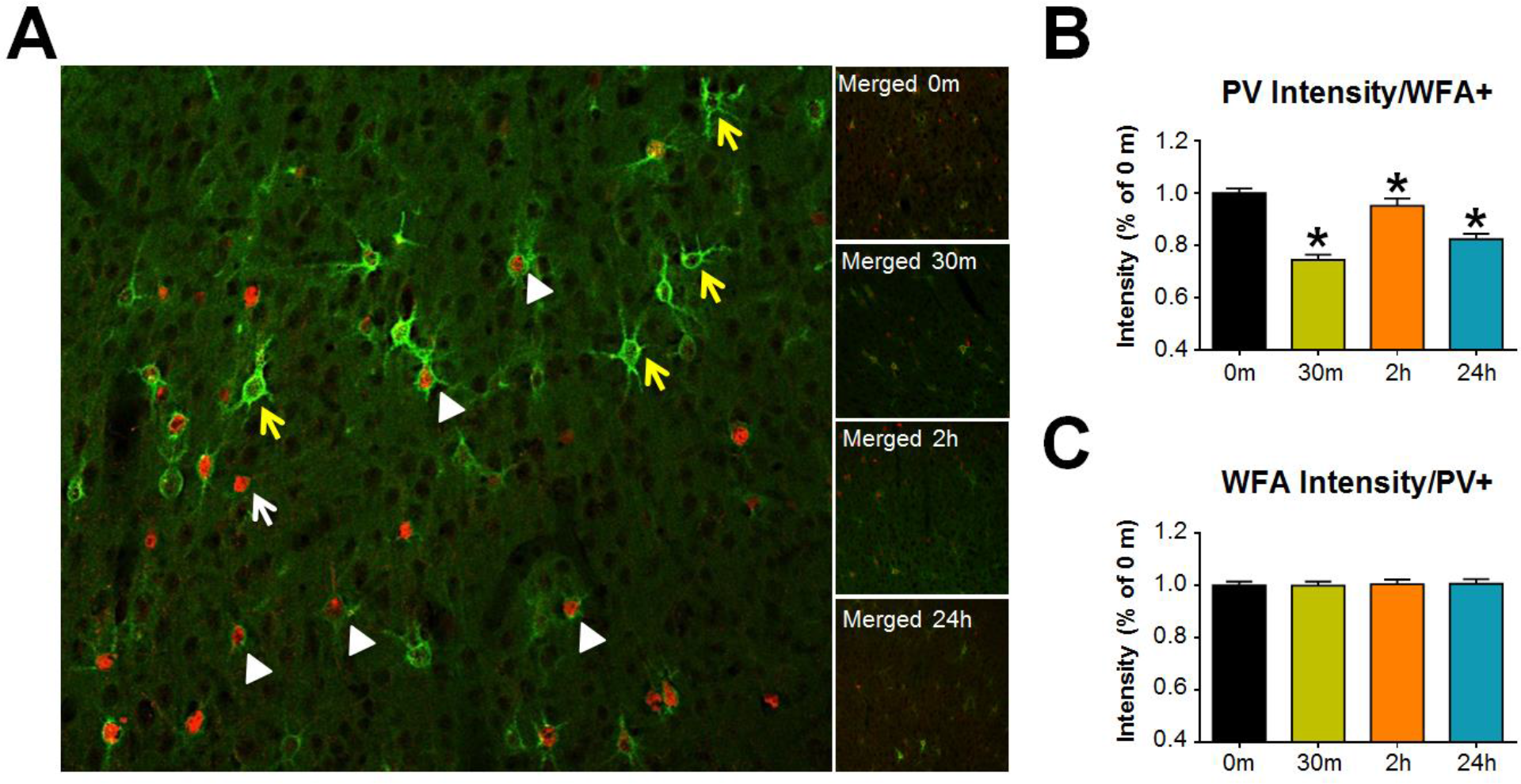
Cocaine memory reactivation decreases PV intensity in WFA^+^− surrounded neurons within the PL PFC. **(A)** Representative photomicrograph of PV+cells with (white arrows) and without (yellow arrows) WFA+ perineuronal nets in the prelimbic PFC. White arrow: PV cells without WFA+ nets; yellow arrows: WFA nets around non-PV cells; white arrowheads: PV+ cells surrounded by WFA+ nets. Left panels are representative images for each time point. **(B)** PV intensity in PV cells surrounded by WFA cells was decreased at all time points after cocaine memory reactivation; **(C)** WFA intensity surrounding PV+ cells were not altered. N-sizes: (t=0 (12); t= 30 m (6); t = 2 h (5); t = 24 h (7)). * P < 0.05, compared with t=0 time point.

### Cocaine-conditioned place preference alters GAD 65/67 and VGluT1 on PV FSIs

We examined excitatory and inhibitory puncta apposing PV cells that were surrounded by PNNs at 2 h and 24 h after memory reactivation. Figure 4A shows a representative PV neuron surrounded by a WFA-labeled PNN, which receives appositions from glutamatergic (VGluT1, magenta) and GABAergic (GAD 65/67, green) puncta (Figure 4B). The number of GAD 65/67 puncta did not change after memory reactivation (Figure 4C), while the number of VGluT1 puncta increased 24 h after memory reactivation compared to t = 0 (Figure 4D; p = 0.02). The increase in VGluT1 and the slight but non-significant decrease in GAD 65/67 led to a decrease in the GAD 65/67:VGluT1 ratio at 24 h compared to t = 0 (Figure 4E; p = 0.02). Figure 4F shows an increase in PV cell diameter at both 2 h and 24 h following cocaine memory reactivation when compared to t = 0 (p<0.0001 for both groups).

**Figure 4.**
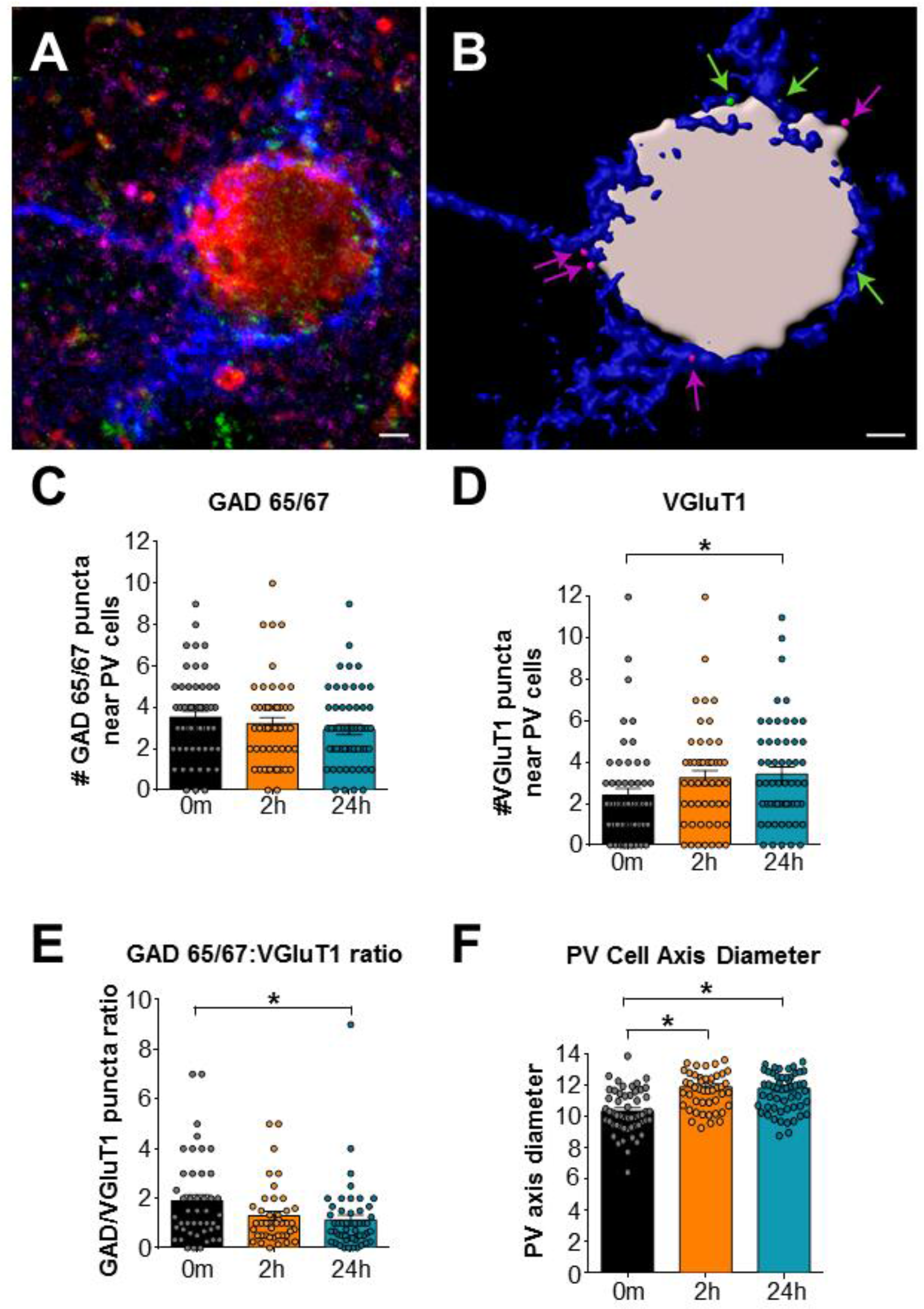
Cocaine memory reactivation alters intensity of glutamatergic (VGluT1) puncta near PNN-surrounded PV neurons and increases PV cell axis diameter. Neurons were visualized with confocal microscopy and analyzed using Imaris segmentation tools. **(A)** Representative PV neuron (red) is surrounded by a PNN labeled with WFA (blue) and receives appositions from glutamatergic (VGluT1, magenta) and GABAergic (GAD65/67, green) puncta. **(B)** The PV neuron (gray) and WFA-labeled PNN (blue) were rendered using the Imaris Surfaces segmentation tool. GAD65/67 (green arrows) and VGluT1 puncta (magenta arrows) meeting our size and location criteria were segmented using the Imaris Spots tool and included in the analysis. Scale bar = 2 μm. **(C)** GAD 65/67 puncta at t = 0 and 2 h and 24 h later; **(D)** VGluT1 puncta at t = 0 and 2 h and 24 h later; **(E)** the ratio of GAD65/67:VGluT1 at t = 0 and 2 h and 24 h later; **(F)** Axis diameter of PV/WFA cells at t = 0 and 2 h and 24 h later. N = 4/group. *P < 0.05, compared to t = 0 time point.

## Discussion

Our previous study showed that modulation of PNN-surrounded neurons (presumably PV+) following memory reactivation reduced reinstatement for cocaine-CPP and increased the activity of pyramidal neurons [21]. In addition, we saw a decrease in the inhibitory tone onto pyramidal neurons but did not identify the underlying mechanism that may have contributed to the increase in pyramidal excitability [21]. This study expands upon our previous work to show that cocaine memory reactivation alters the electrophysiological properties of PNN surrounded FSIs (Figures 1 and 2), and corresponds with a decrease in PV intensity (Figure 3B) within the PL PFC, suggesting a mechanism by which inhibitory tone onto pyramidal neurons may be reduced to increase network excitability and ultimately promote cocaine-seeking behavior.

PV is a calcium-binding protein found primarily within FSIs and plays a role in modulating the electrical properties of FSIs by influencing neuronal excitability and activity [11, 44]. Within the PFC, PV FSIs preferentially inhibit pyramidal neurons and refine pyramidal neuron output and timing, which influences learning and memory [14]. For instance, Pavlovian contextual fear conditioning and environmental enrichment differentially affect PV and GAD67 expression to influence network plasticity [19]. Moreover, in PV knockout mice, hippocampus-dependent tasks are impaired, and AMPAR-mediated currents are decreased [45]. Finally, schizophrenic patients have a reduction in PV neurons, which may contribute to cortical network dysfunction [46, 47].

Mechanistically, in addition to PV’s ability to bind calcium, PV has also been implicated in the recruitment and expression of potassium channel subunits Kv3.1 and Kv3.4 as well as the calcium channel Cav3.2 [48–51]. This specific potassium channel subunit composition is almost exclusively expressed within PV neurons and changes when the level of PV decreases [50]. PV levels are also inversely related to cell and mitochondria volume and reactive oxygen species (ROS) production [52–55], suggesting that the conditioned effects of cocaine-associated memory reactivation could decrease PV through ROS production. Additionally, GAD 67 levels are correlated with NMDAR subunit composition [56]. Specifically, lower levels of PV resulted in less GABA release and a decrease in inhibition. This may explain why we observed a reduction in PNN-surrounded PV FSI excitability (Figure 1F) following cocaine memory reactivation. Moreover, the increase in inter spike interval (ISI) that we observed 30 m and 2 h (strong trend but not significant) after reactivation could account for the attenuation in FSI excitability (Figure 1G). Additionally, we found an increase in glutamatergic synaptic input onto PNN surrounded PV FSIs, as supported by our mEPSC data 30 min, 2 h, and 24 h after memory reactivation (Figures 2A & 2C). Although we did not see a significant change in the average frequency and amplitude of mIPSC events, there was an overall trend towards a reduction in inhibitory transmission, which was significant when analyzing the cumulative data of all the miniature events (Figure 2F & 2H). This increase in excitatory:inhibitory balance onto PNN surrounded PV FSIs was corroborated by the GAD65/67:VLGut1 ratio (Figure 1F). We speculate that this increase in excitatory neurotransmission may be a compensatory response as previously reported following prolonged withdrawal from cocaine [57]. Moreover, we saw changes in both frequency and amplitude of mEPSC events, indicating that there may be both pre- and post-synaptic changes following memory reactivation, when the memory is in a labile state [58]. One proposed explanation for the change in excitatory:inhibitory balance onto PNN surrounded PV FSIs could be that as inhibition of PV cells onto local pyramidal neurons is decreased, those pyramidal neurons become more excitable back onto PV FSIs. Although intrinsic excitability returns to levels indistinguishable from controls at 24 h, excitatory synaptic tone remains elevated, suggesting that a more complex mechanism is involved when the window of plasticity in memory reconsolidation closes. An alternative explanation could be that there are other components of the tetrapartite synapse being altered following cocaine memory reactivation that we did not directly measure. For example, some PNN components exist not only in the ornate lattice structure surrounding the neuron, but also within the general extracellular matrix [62,63]. Although we did not find differences in WFA intensity (Figure 3C) on PNN surrounded cells, there could be differences in the ECM or PNN components (such as aggrecan, a main component of PNNs) that we have not yet identified.

PNNs are modified by a variety of stimuli, including cocaine exposure [34, 59], cocaine-CPP acquisition [21], alcohol [60, 61], nicotine self-administration [62], heroin self-administration [63], sleep deprivation [40],environmental enrichment [64], and consumption of high fat diets [32]. A majority of PV FSIs within the PL PFC are surrounded by PNNs, and previous work by our group has shown that the amount of cocaine exposure differentially affected PNN and PV expression [34]. Specifically, we showed that 1 d of cocaine exposure decreased WFA intensity (a marker for PNNs) but 5 d of cocaine exposure increased intensity [34]. In addition, we showed that PV intensity changed similarly to WFA intensity but generally lagged in the timing of the change. However, here we found no changes in WFA intensity following cocaine-associated memory reactivation (Figure 1C). This was surprising to us since PV and PNNs tend to fluctuate and develop in synchrony, working together to modulate plasticity and cortical development [65]. Yet, there are a couple instances where this is not the case. For instance, Harkness and colleagues found that PV levels increased after 12 h of sleep deprivation while PNN intensity remained unchanged [40]. In addition, within the hippocampus, PV and PNN development and expression do not coincide, which is different from what is seen in cortical areas [66]. Hence, although changes in both PV and PNNs typically have a high degree of correlation, there are circumstances when this does not hold up, and cocaine-CPP memory reactivation is another example of this divergence. We also cannot rule out the possibility that there could be a failure in PNN stability/formation and consequently WFA staining is impaired due to an inability for the WFA to bind [67]. Future studies could use selective antibodies for specific proteoglycans to potentially rule this out. Our data suggest differential roles for PNNs and PV in the formation and maintenance of cocaine-associated memories. Perhaps PNNs are more important for the initial encoding of cocaine associations and PV is more involved in memory maintenance after the memory has been established.

In conclusion, we show for the first time that cocaine-associated memory reactivation reduces excitation of PNN surrounded FSIs (Figure 1F), increases the excitatory:inhibitory balance onto these neurons (Figures 2 & 4), and decreases PV intensity (Figure 3B). These adaptations in electrophysiological function may account for the changes we previously saw in pyramidal neuron firing [21]. In addition, we saw a divergence in PV and PNN expression following memory reactivation, which is different from what we found after cocaine exposure [34]. This suggests that PV may play a role in memory reactivation independent from PNNs. Whereas PV may be necessary for memory stability and recoding, PNNs may be necessary during drug acquisition and modification following cocaine exposure to recapitulate the developmental process [68]. Future studies are examining the underlying mechanistic changes that mediate adaptations in PNN-surrounded PV FSI excitability. Specifically, future experiments will work to characterize the role of potassium and/or calcium channel expression,distribution, and mediated currents following memory reactivation. Taken together, this work established that cocaine-associated memory reactivation alters cortical network function *via* modification of PV neurons, which would impact PFC pyramidal cell output and in turn drug-seeking behavior.

## Acknowledgments

The authors would like to thank Dr. Paige Dingess for her help and support of this work. This work was supported by the National Institute on Drug Abuse R01 DA040965 (TEB and BAS), the National Institutes of Health Centers Program Grant P30 GM 103398-32128 (TEB), National Institute of Neurological Disorders and Stroke P30 NS061800 (SAA), the Wyoming Scholars Program (DJB), and the University of Wyoming Science Initiative (DJB).

## Authors Contribution

TEB and BAS were responsible for the study concept and design. ETJ, JHH, AEG, DJB, and JAA contributed to the acquisition of animal behavior. ETJ drafted the manuscript, performed all electrophysiology, analyzed and interpreted electrophysiological data. JHH and AEG performed immunohistochemical procedures, collected, and analyzed images. DMH performed all puncta data collection and analysis. TEB, BAS, and SAA provided critical revision of the manuscript. All authors critically reviewed and approved final version for publication.

